# A survey of fishes of Hombolo Lake, Dodoma, Tanzania, with evidence for local extinction of a native tilapia as a consequence of stocking

**DOI:** 10.1101/452847

**Authors:** George F. Turner, Benjamin P. Ngatunga, Martin J Genner

## Abstract

The fish community of the Hombolo Lake, an impoundment on the Wami catchment near Dodoma, Tanzania, was surveyed in 2014 and 2017. The lake contains a relatively low diversity community dominated by two *Oreochromis* species introduced from outside the Wami catchment, *O. niloticus* and *O. esculentus*. Evidence from historical collections suggests that the native *O. urolepis* was formerly present, and its current absence is likely to be the result of competitive exclusion or genetic swamping by non-native species introduced for fishery enhancement. Four other fish species were also recorded.

## Introduction

Tanzania is a global hotspot of tropical fish biodiversity and, in particular, houses the largest diversity of native tilapia species of the most important farmed genus *Oreochromis* of any country (Lind *et al*. 2012). Recent studies have shown that stocking of exotic strains of tilapia can have detrimental effects on native species, through competitive exclusion and genetic swamping following hybridisation (Angienda et al. 2011; D’Amato *et al*. 2007, Deins *et al*. 2014, Ndiwa *et al*. 2014). Until recently, little information has been available about the current distributions of native and exotic species in Tanzania. Here, we focus on Lake Hombolo, a small (7x 1.4 km) water body in central Tanzania, located in the Dodoma region; S05°57’02.8”; E035°58’07.6” at an elevation of approximately 1040m (Figures 1-4). It is an impoundment of a small tributary stream on the Upper Wami River catchment, reportedly in the 1957 (Petr 1974), originally for the flood control, irrigation and fisheries. According to local officials, it is presently used to irrigate vinyards. The water is rather saline and alkaline, with conductivities of 3,300-4,000 μScm^−1^ and pH of 8-8.5 (Shemsanga *et al*. 2017). In 1972-73, values of 1650-2,400 μScm^−1^ and pH 7.8-8.1 were obtained, suggesting increasing salinity over time (Petr 1974). At that time, maximum water depth was reported as being 12m. An unpublished report on the fish fauna (Petr 1974), reported the following species (classification here updated) in the lake during surveys in 1972-73: *Astatotilapia bloyeti, Clarias gariepinus, Coptodon zillii, Enteromius paludinosus, Labeo cylindricus, Oreochromis esculentus, Oreochromis urolepis*. Voucher specimens of cichlid species collected by the author were deposited in the collection of the Natural History Museum in London, and we examined the specimens of *O*. urolepis.

**Figure 1:**
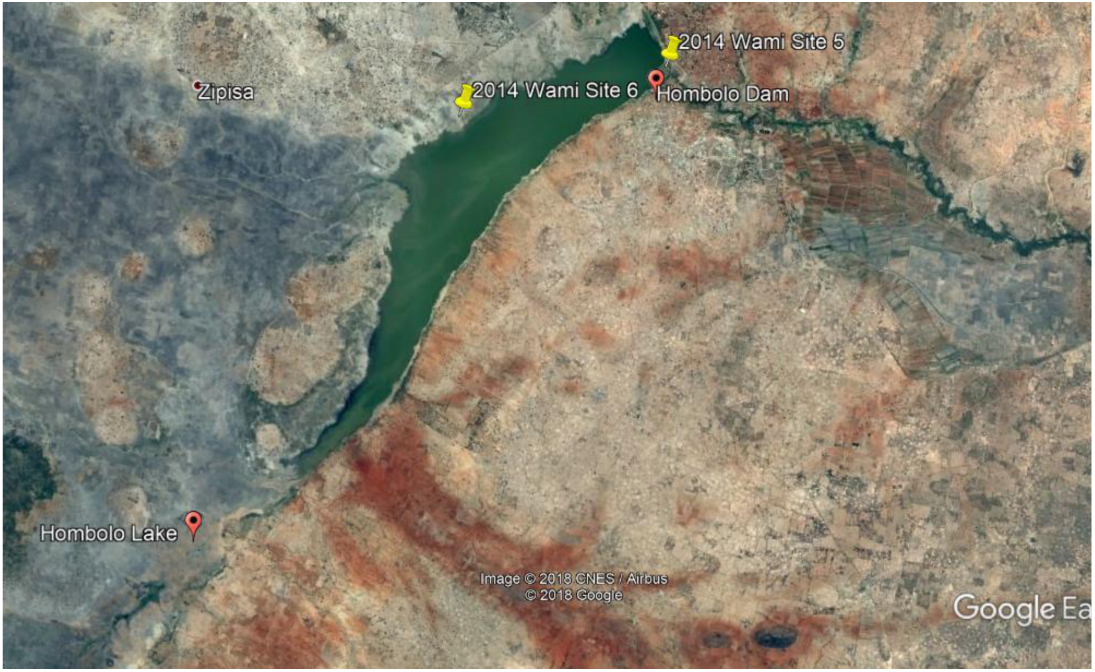
Hombolo Lake. Google Earth

**Figure 2:**
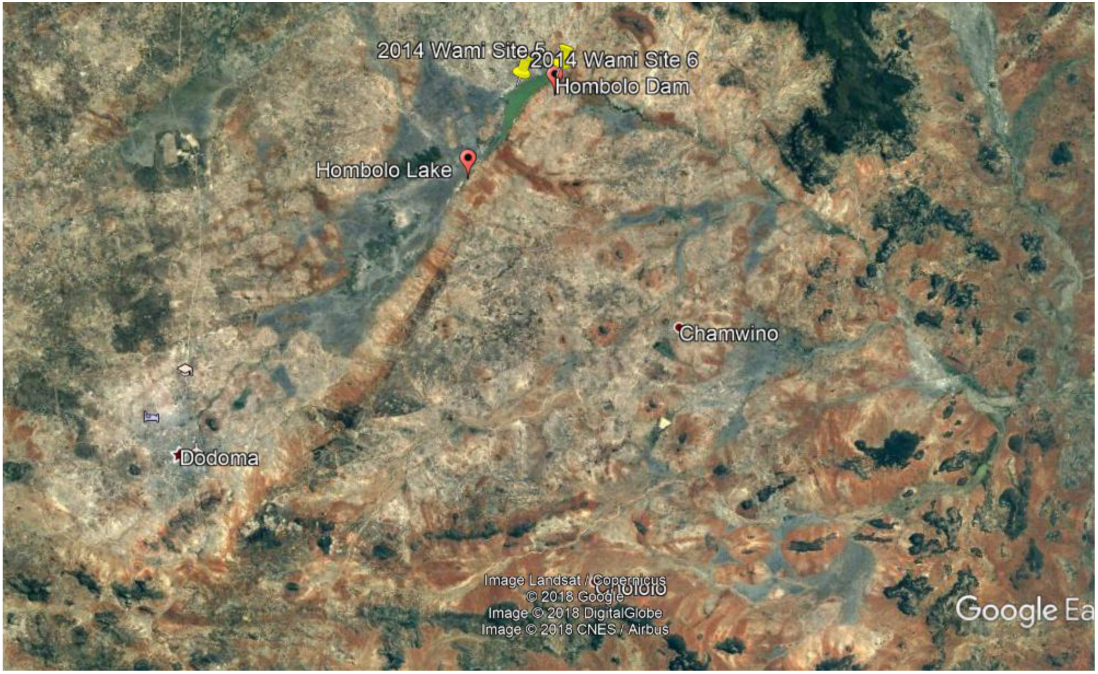
Location: ca 28km NE of Dodoma. Google Earth

**Figure 3:**
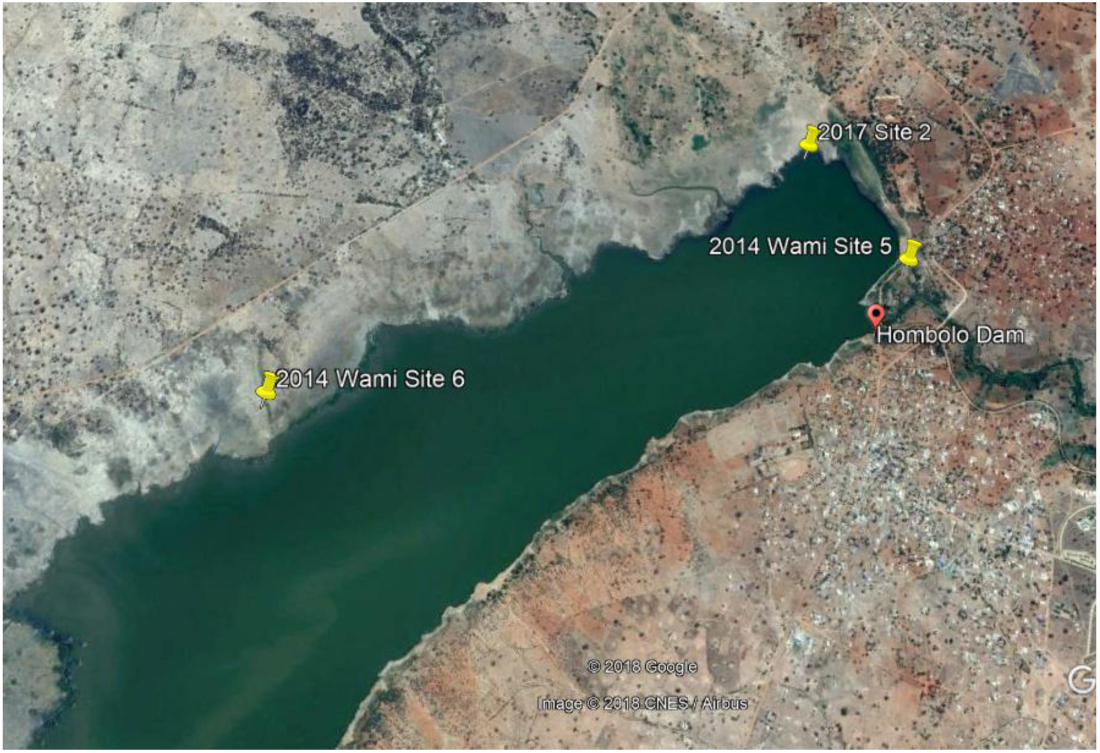
Hombolo Lake Sampling Sites: NB Site 1 in 2017 was the same as Site 5 in 2014. Google Earth

**Figure 4:**
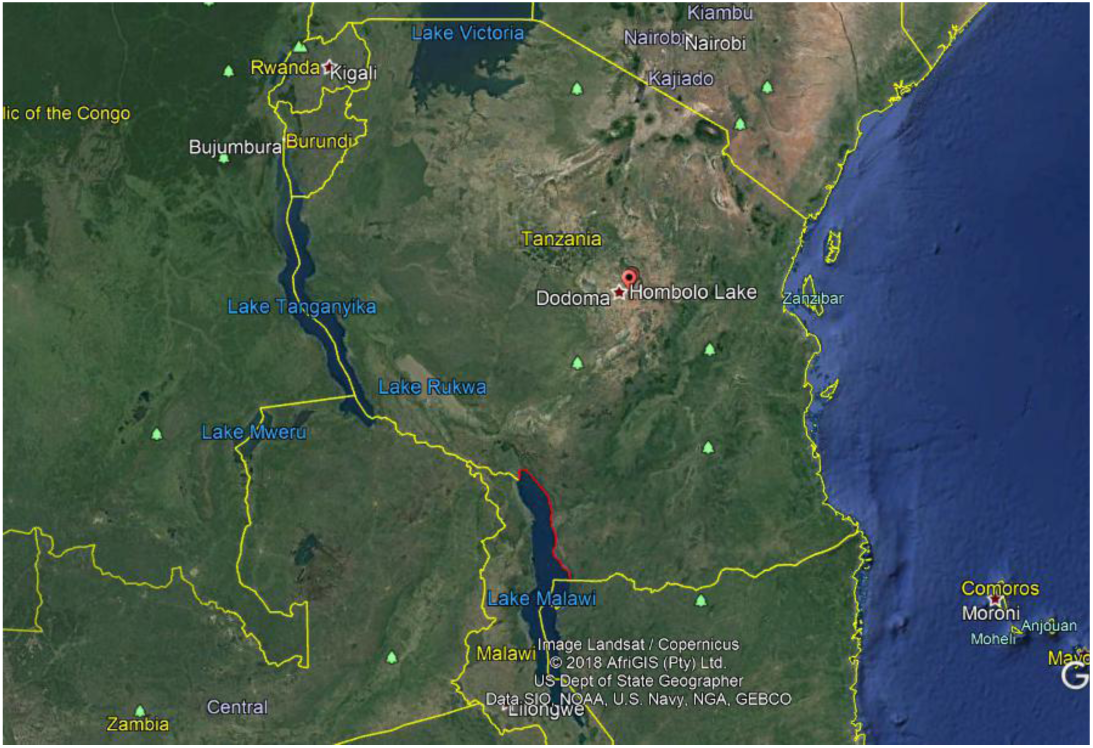
Location of Hombolo Lake within Tanzania. Google Earth

## Methods

The lake was sampled during January 2014 (BPN, GFT) and January 2017 (GFT, A.G.P. Ford, A.Shechonge, A.M. Tyers). Specimen identifications were checked with type material in international museum collections. Voucher specimens, photographs and fin clips for DNA analysis have been archived at respective institutions.

## Results

When visited, the water was very turbid, with some remaining standing dead trees and fringed with banks of reeds/papyrus. At the shoreline sites, the water remained very shallow for a long distance offshore. Active fisheries with gillnets, beach seines and mosquito seines were seen (Figures 5,6).

**Figure 5:**
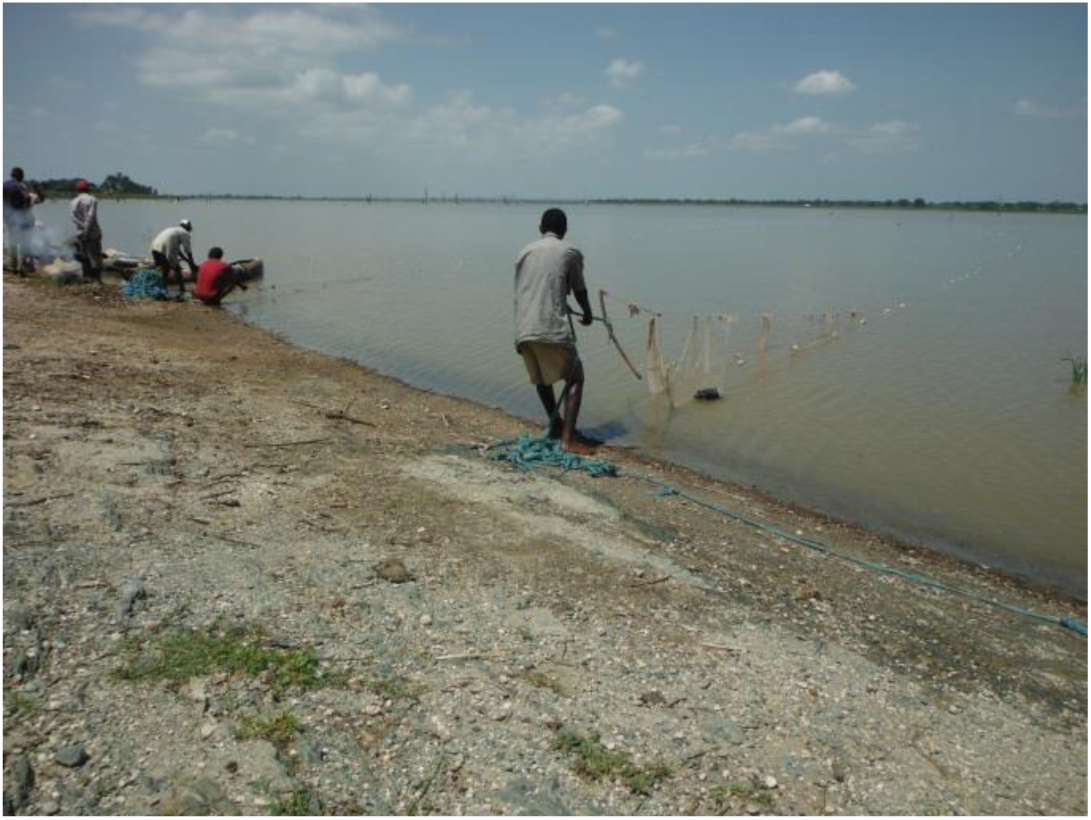
Hombolo Lake, 2014: Seine netting

**Figure 6:**
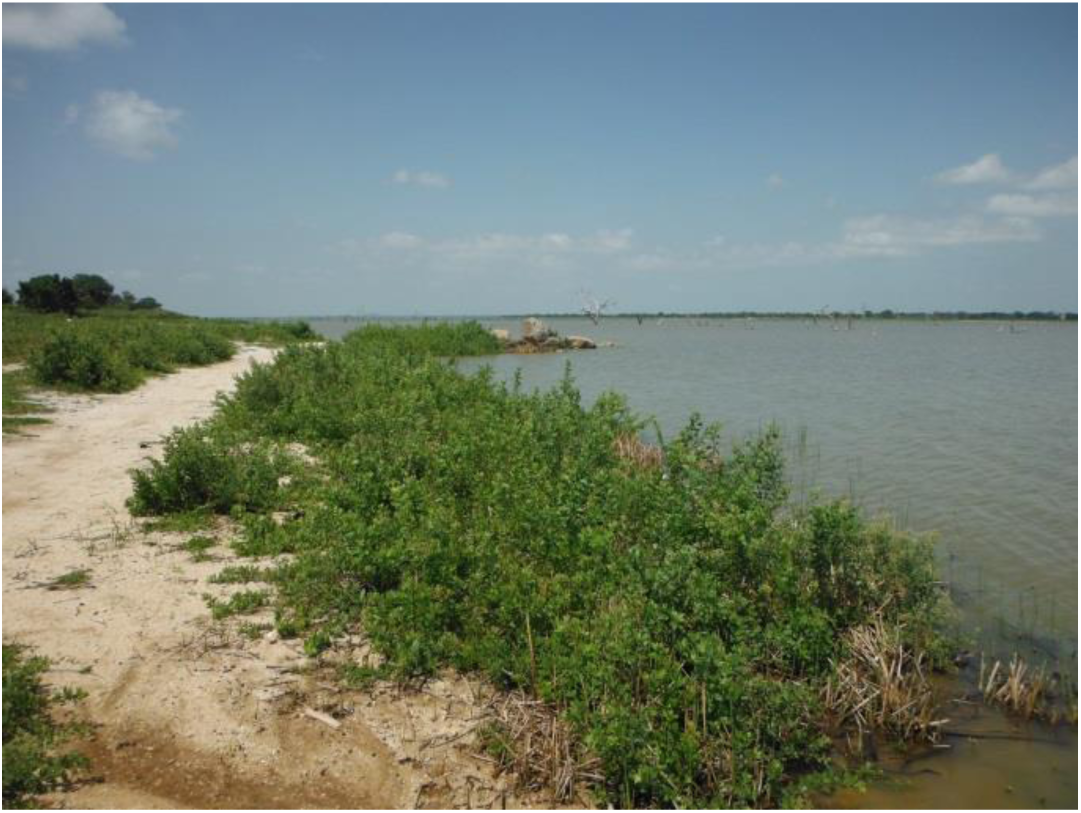
Hombolo Lake, 2014

**2014 Sampling:** 28^th^ January 2014, 2014 Wami Site 5: Hombolo Dam, near Dam wall. S05°57’02.8”; E035°58’07.6”. Elevation 1037m. Samples bought from gillnetters and seine netters included the dominant *Oreochromis niloticus* (L. 1757) (Figure 7), a few *Oreochromis esculentus* (Graham 1928) (Figure 9) and a single *Enteromius cf. paludinosus* (Peters 1852) (Figure 10).

**Figure 7:**
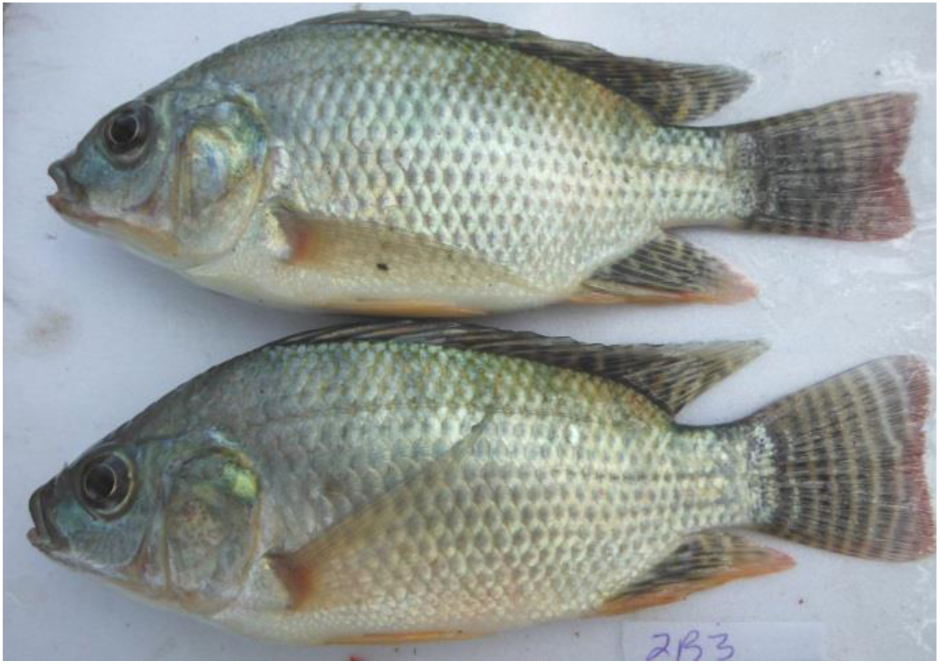
*Oreochromis niloticus*, Hombolo Lake 2014

**Figure 9:**
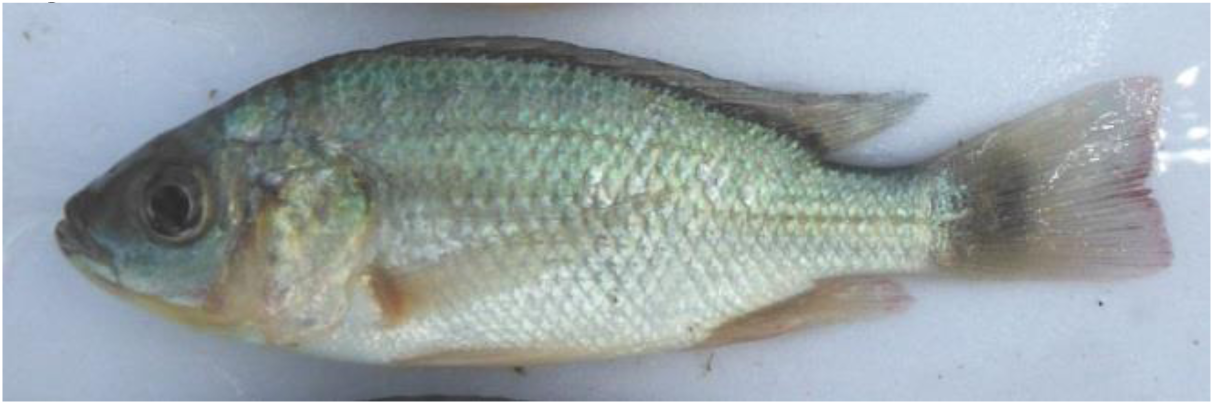
*Oreochromis esculentus*, Hombolo Lake 2014

**Figure 10:**
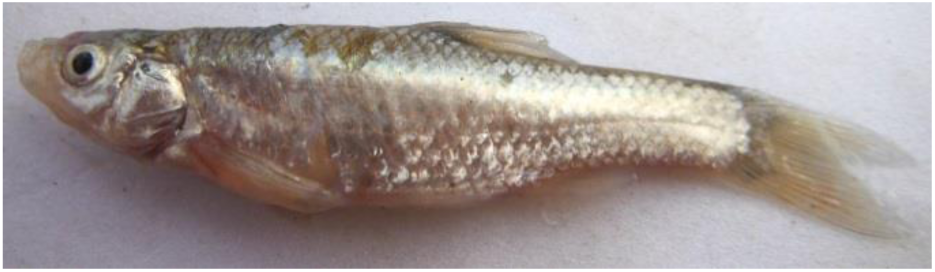
*Enteromius cf. paludinosus*, Hombolo Lake 2014

2014 Wami Site 6: Hombolo Lake. S05°57’08.2”; E035°56’34.6”. This site was accessible by driving and then walking across the sun-baked mud of the dry lake bed. Samples were bought from seine netters camped at the shoreline several hundred metres from the permanent bank. All specimens seen here were *Oreochromis esculentus*, including large numbers of relatively small males in breeding dress (Figure 8), suggesting that there was a breeding aggregation nearby.

**Figure 8:**
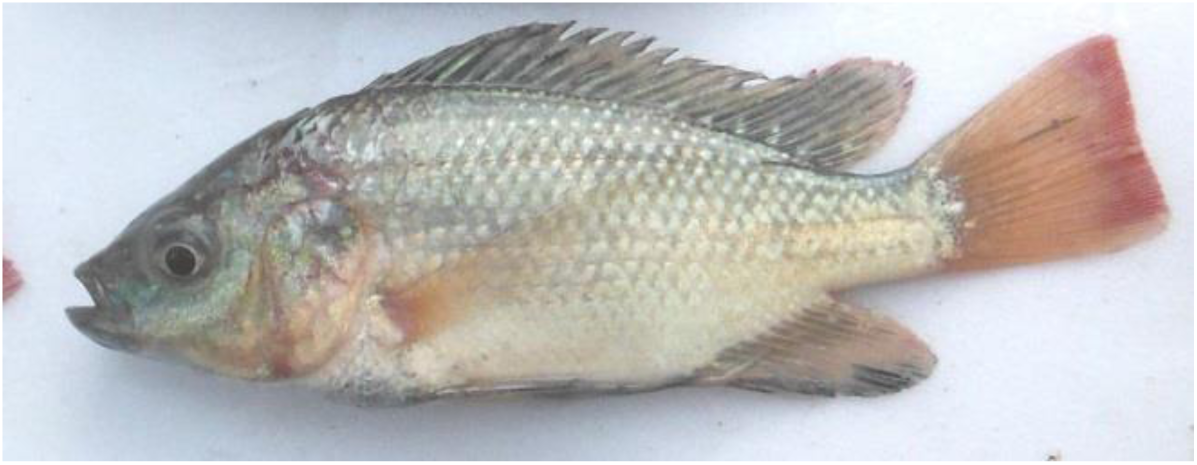
*Oreochromis esculentus*, Male Hombolo Lake 2014

**2017 Sampling:** 10^th^ January 2017. 2017 Wami Site 1: We revisited the dam wall (2014 Wami Site 5): local gillnetters had landed a catch already (Figure 11). The gillnet catch was again dominated by *O. niloticus* (Figure 13), with a few *Clarias cf. gariepinus* (Burchell 1822) also seen. There was no sign of *O. esculentus* being landed. The area to the north of Site 1 was very muddy and judged impassible by car. We approached the lake on foot (at 2017 Site 2: Figure 12), but found no fishing in progress apart from 2 people who seemed to be catching shrimps with some kind of makeshift net. We hired them to use a length of mosquito netting as a seine in shallow water. This yielded a large number of *Astatotilapia bloyeti* (Sauvage 1883) (Figure 15), including many rather small males showing both blue and yellow background colour. Other species landed included a few *Enteromius cf. paludinosus* and a single *Coptodon cf rendalli* (Boulenger 1897) (Figure 14). With the assistance of a local official, we were able to obtain 3 specimens of *O. esculentus* from a trader, from an unknown location on the lake. It was not clear whether the relative rarity of this species was due to declining populations or targetting of the fisheries in operation at the time of visiting.

**Figure 11:**
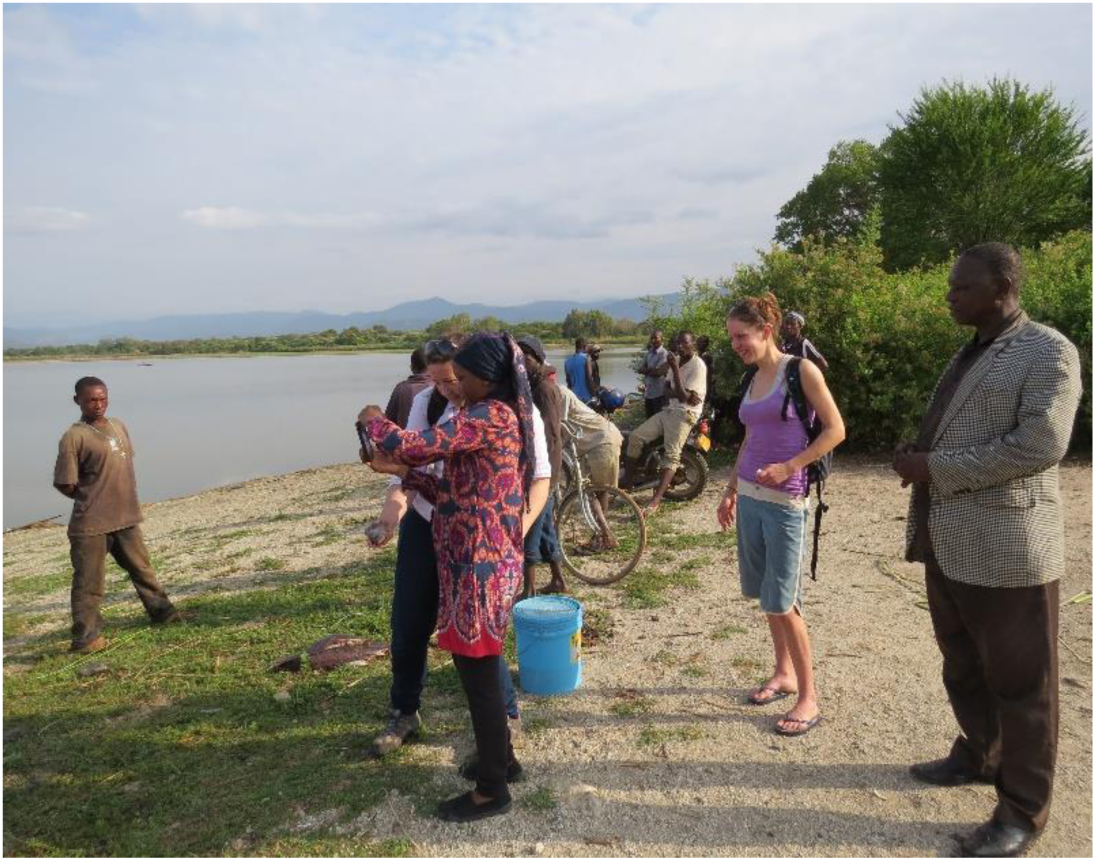
Hombolo Dam 2017 Site 1.

**Figure 12:**
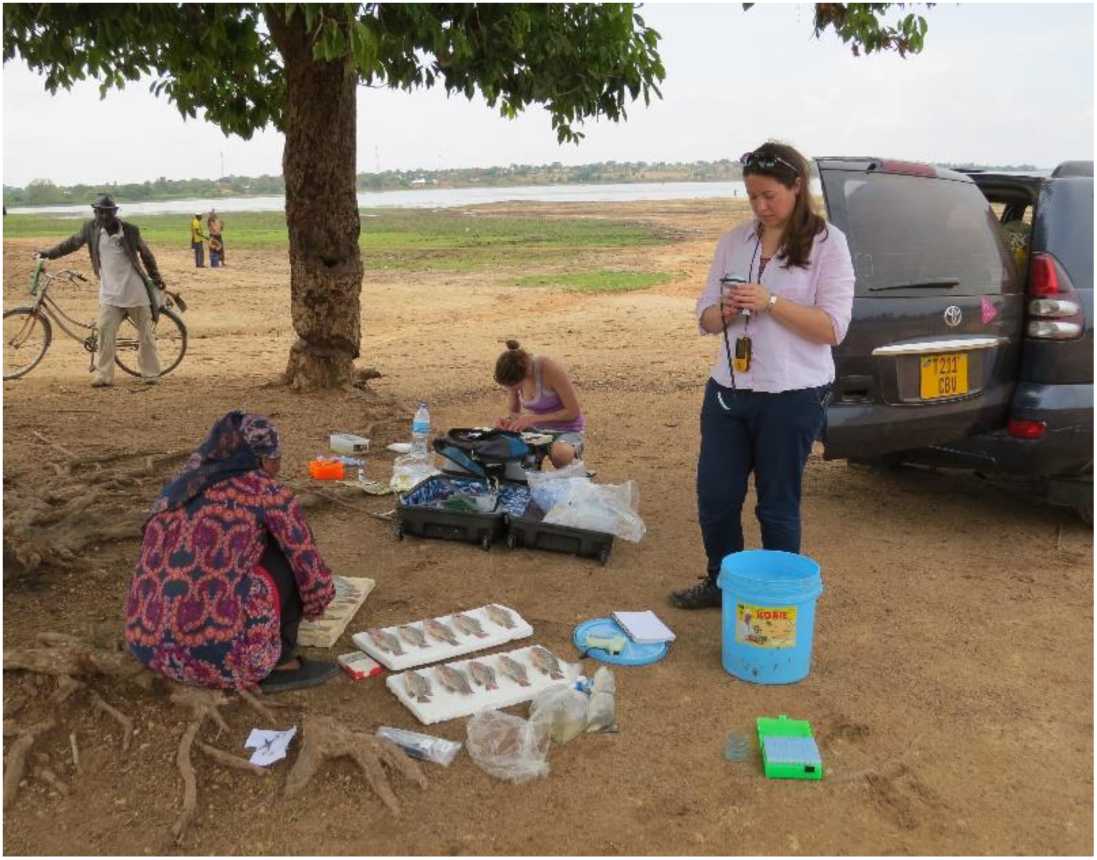
Hombolo Lakeshore sampling 2017 Site 2

**Figure 13:**
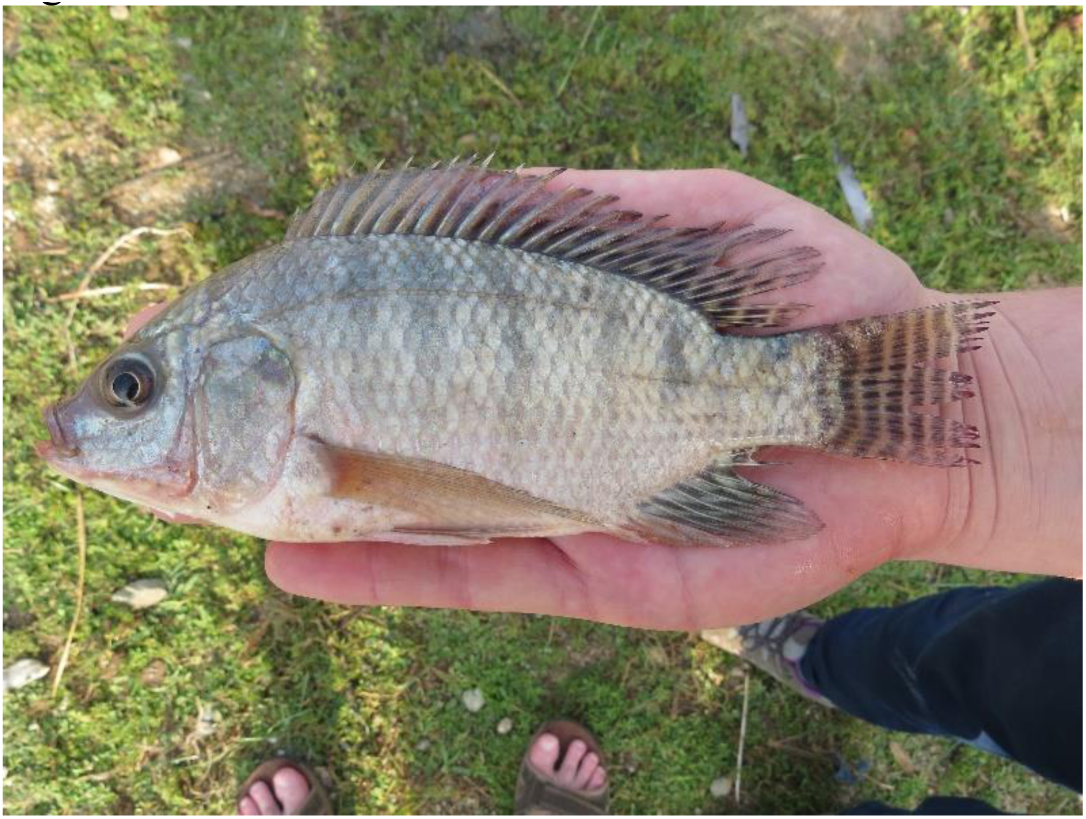
*Oreochromis niloticus*, Hombolo Lake 2017

**Figure 14:**
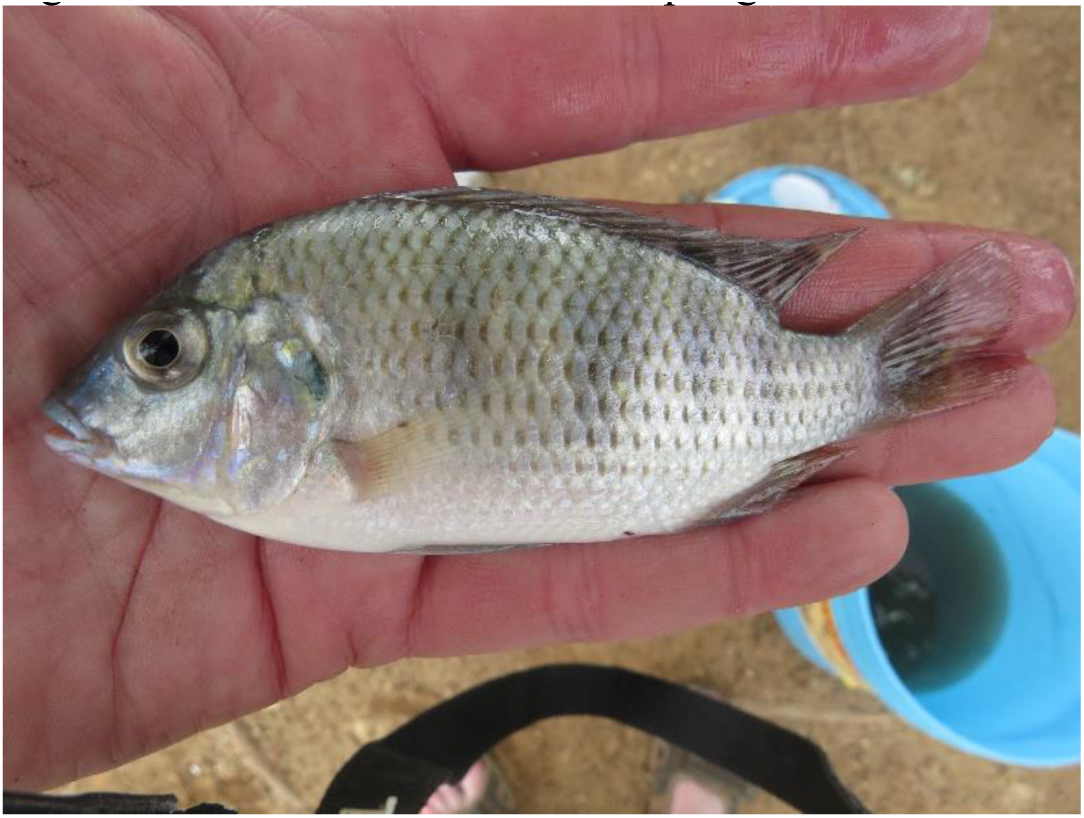
*Coptodon cf. rendalli*, Hombolo Lake 2017

**Figure 15:**
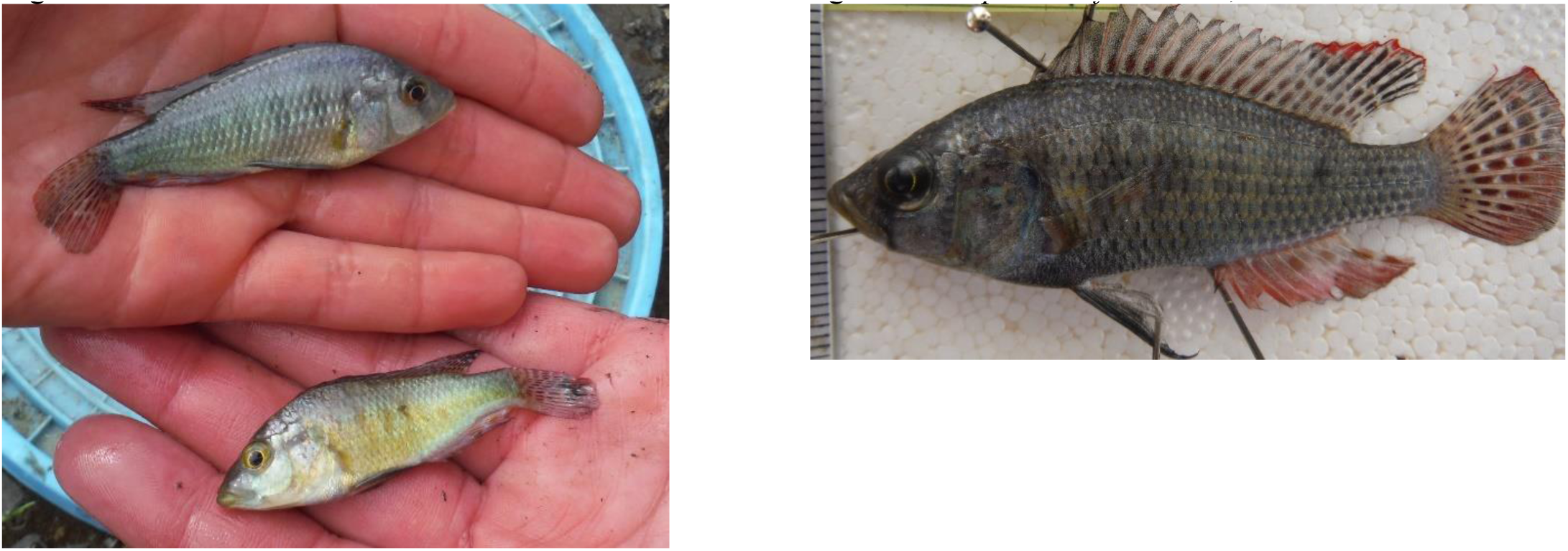
*Astatotilapia bloyeti*, Hombolo Lake, 2017. Freshly collected males varied considerably in background colour (left). A pinned specimen illustrates the characteristic fin markings of adult males (above)

**Specimens from Natural History Museum in London:** During a visit to the NHM on 8-9 February 2016, specimens of *Oreochromis* from Homobolo Dam were examined by GFT. These were catalogued as BMNH 1973.5.21: 217-223: *Tilapia hornorum*, Hombolo Dam, Upper Wami System, 17/6/72-13/6/72 (sic), coll T. Petra (presumably Petr), pres. R. Bailey. The material comprised seven small specimens, 97, 69, 62, 60, 58, 57 & 56 mm SL. The smallest three were badly bent. All specimens were pale coloured with conspicuous thin dark bars and a ‘tilapia mark’ on the dorsal fin. In some specimens, the caudal fins had a few prominent stripes. They were photographed but not measured (Figure 16 A-D).

**Figure 16:**
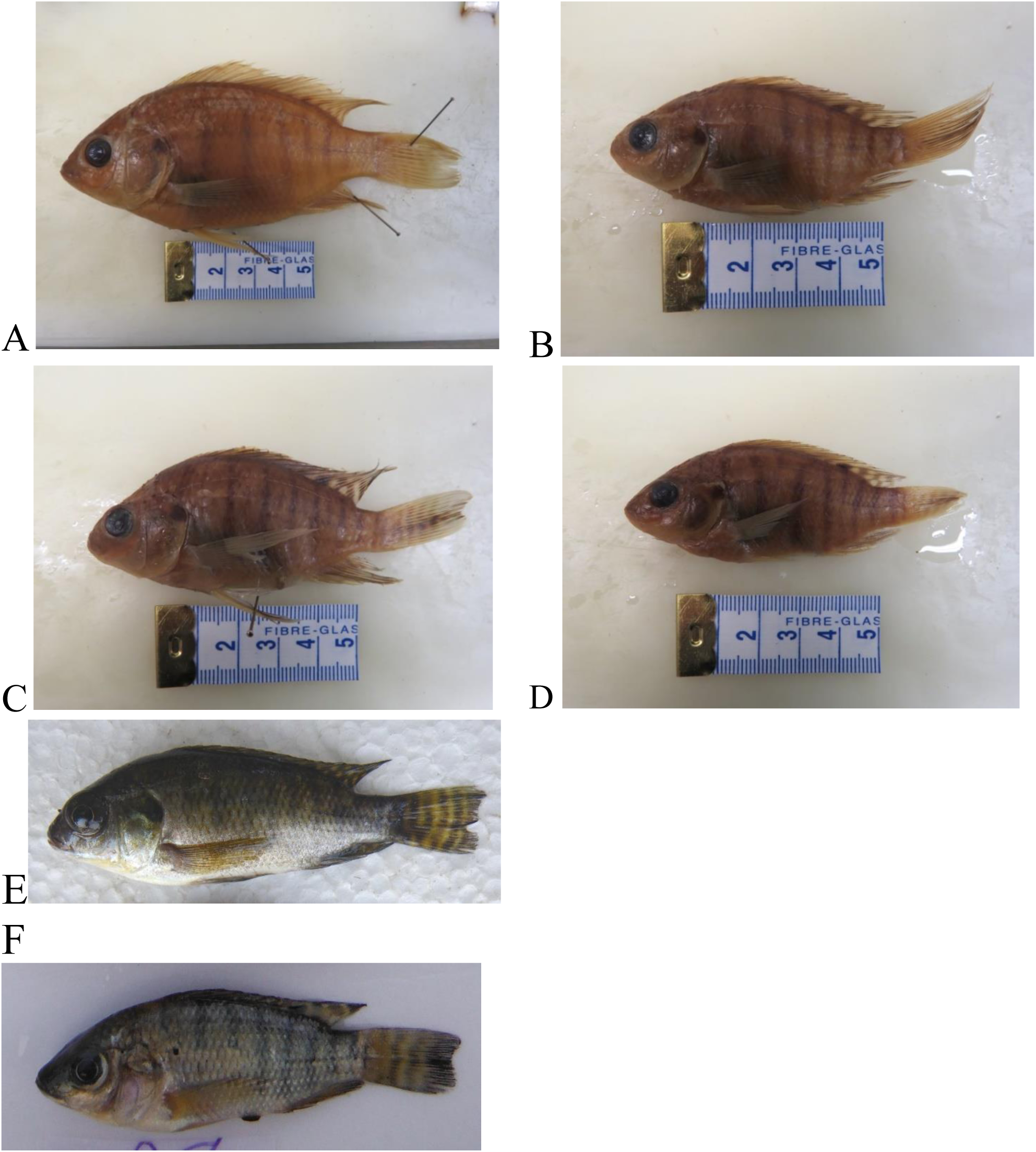
A-D Specimens of *Oreochromis* sp in London, Natural History Museum, collected from Hombolo Dam in 1972, labelled as ‘*Tilapia hornorum*’ E. Small indvidual of *O. niloticus* collected from Lake Itamba (Mbeya District) in 2011. F. Small individual of *O. urolepis* collected from the lower Wami River in 2014. Specimen C shows jagged, more vertical stripes, that are fainter nearer the caudal peduncle, which fits better with the fresh *O. urolepis* specimen (F) than the *O. niloticus* specimen (E).

The London specimens (Fig 16A-D) appear to be *Oreochromis urolepis* (Norman 1922), a species native to the Wami system, but not recorded in the 2014 or 2017 sampling. They certainly are not *Oreochromis esculentus* (Figs. 8,9), and Petr’s (1974) report indicates that both species were present and were distinguished in the field. According to Trewavas (1983, page 211), diagnostic features of *O. esculentus* include a slender caudal peduncle and absence of stripes on the caudal fin, which are consistent with the 2014 specimens illustrated above, but not with these 1972 specimens. The other *Oreochromis* species presently found in the lake is *O. niloticus*, which also has striped caudal fins (Figure 7, 13). There are no clearly diagnostic morphological differences between this species and *O. urolepis* at such small sizes (Trewavas 1983). However, *O. niloticus* usually has more consistently prominent and regular stripes in the caudal fin than *O. urolepis*. Comparison of similarly-sized strongly-striped individuals of the two species (e.g Figure 16 E-F) indicates that the stripes of *O. niloticus* tend to be regular, gently curved and are appear dark close to the caudal peduncle, while those of *O. urolepis* tend to be straight, but rather jagged and fade closer to the body. The London specimens also have rather deep caudal peduncles, which is characteristic of *O. urolepis*. In any case, it would be surprising if *O. niloticus* was in Hombolo Lake in 1972: it was not reported by Petr, who at that time gave his address as Makere University in Uganda, a country where Nile Tilapia is native and widely exploited. Nile Tilapia was not widely stocked in Tanzania in the 1970s and when interviewed in 2014, local fishermen claimed *O. esculentus* to be native but *O. niloticus* to have been accidentally introduced in the 1990s when flooding led to overflow of nearby aquaculture ponds. Thus, it seems likely that the 1972 specimens are correctly identified as ‘*Tilapia hornorum* Trewavas 1966’, a junior synonym of *O. urolepis. Oreochromis urolepis* is native to the Wami system, and *O. hornorum* (sometimes considered a distinct subspecies) was originally described from specimens collected on this system near Kilosa on the Mkondowa River. Thus, it appears that the lake was almost certainly originally populated by the native *Oreochromis urolepis*.

## Discussion

At the time of sampling in 2014-17, the fish fauna of the lake was relatively depauperate, with only 6 species recorded. Of these, three have almost certainly been introduced: *Oreochromis niloticus, O. esculentus* and *Coptodon cf. rendalli*. Petr (1974) reported that the major fishery species (‘tilapia’ and Labeo) were introduced, because there were few fish species in the inflowing streams. Within Tanzania, the Nile Tilapia, *O. niloticus*, is native only to Lake Tanganyika and its catchment, but it was stocked into Lake Victoria in the 1950s, initially probably from Lake Albert in Uganda (Trewavas 1983), perhaps supplemented by stocks from elsewhere. The Nile Tilapia has been collected from Lake Victoria for production in hatcheries and distribution to fish farmers throughout Tanzania since the 1990s. It has also been widely stocked in natural lakes and impoundments of natural river systems (Shechonge *et al*. 2018a). *Oreochromis esculentus* is endemic to the catchment of Lake Victoria (Trewavas 1983), where it has largely been replaced by *O. niloticus* (Shechonge *et al*. 2018a). It was widely stocked in Tanzania during the 1950s and 60s, most notably in Lakes Singida, Nyumba ya Mungu (Trewavas 1983) and Rukwa (Seegers 1996). Our surveys since 2011 have indicated its continued occurrence in these lakes (Shechonge *et al*. 2018a). *Coptodon rendalli* is native to Tanzania, occurring naturally in the catchments of Lakes Tanganyika (Eccles 1992) and Malawi (Nyasa), the type locality being the Lake Malawi outflow, the Upper Shire River (Boulenger 1897). Eccles reports that it has been widely distributed through stocking and fish farming. In our recent surveys, *Coptodon* was found in almost all water bodies surveyed. Identification is slightly complicated by the fact that *Coptodon rendalli* and the very similar-looking *C. zillii* (Gervase 1848) (not native to Tanzania) have both been stocked in Lake Victoria. Petr recorded ‘Tilapia zillii’ in Hombolo in the 1970s. *Coptodon* have been seen in hatcheries in Tanzania where it was claimed that Nile Tilapia, originally sourced from Lake Victoria, were being bred for distribution to fish farmers: if the *Coptodon* were also from Lake Victoria, these could be either or both species or indeed hybrids.

Two of the other species, provisionally identified as *Clarias gariepinus* and *Enteromius paludinosus* are widespread and very abundant in most water bodies in the region (Eccles 1992), so it is probable that they are native. Work in progress indicates that *E. paludinosus* represents a complex of species. The final species observed, the small haplochromine cichlid *Astatotilapia bloyeti*, is indigenous to the Wami system (type locality near Kilosa) and to other river systems to the north and west and so is very likely to be naturally occurring. Petr (1974) reported *Labeo cylindricus* to be abundant in experimental gillnet catches, but we did
not record this species at all.

The absence of *Oreochromis urolepis* from our recent collections is noteworthy. As this species is native to the Wami system and appears to have been present in the lake in the 1970s, the likeliest explanation is that it has been exterminated through competitive exclusion and/or genetic swamping by the introduced *O. esculentus* and/or *O. niloticus*. In the 1970s, both *O. esculentus* and *O. urolepis* were abundant in the lake, and so competitive exclusion of the latter by the former seems unlikely. *Oreochromis urolepis* is known to have a higher salinity tolerance than *O. niloticus* (Trewavas 1983), so increased salinization would not be expected to favour the latter. If *O. niloticus* can competitively exclude *O. urolepis*, even in a relatively saline lake, this does not bode well for the long-term survival of the latter species, given that *O. niloticus* is established in all major catchments where *O. urolepis* is native (Shechonge et al. 2018a) and that the two species are known to hybridise (Shechonge *et al.* 2018b).

## Acknowledgements

We are grateful to Antonia Ford, Asilatu Shechonge and Alix Tyers for help with fieldwork, to Oliver Crimmen, Simon Loader and James Maclaine with help accessing the collections at the Natural History Museum (London) and to Royal Society/Leverhulme Trust Africa Awards under the project MolecoFish (to MJG, BPN, GFT) and a BBSRC/NERC Aquaculture grant (to GFT and Federica Di Palma) for financial support, and to TAFIRI and COSTECH for help with logistics and permits.

## References

Angienda PO, Lee HJ, Elmer KR, Abila R, Waindi EN, Meyer A (2011) Genetic structure and gene flow in an endangered native tilapia fish (*Oreochromis esculentus*) compared to invasive Nile tilapia (*Oreochromis niloticus*) in Yala swamp, East Africa. Conservation Genetics 12: 243–255.

Boulenger GA (1897) Descriptions of new fishes from the Upper Shiré River, British Central Africa, collected by Dr. Percy Rendall, and presented to the British Museum by Sir Harry H. Johnston, K. C. B. Proceedings of the Zoological Society of London 1896: 915–920.

D’Amato ME, Esterhuyse MM, Van Der Waal BC, Brink D, Volckaert FA (2007) Hybridization and phylogeography of the Mozambique tilapia *Oreochromis mossambicus* in southern Africa evidenced by mitochondrial and microsatellite DNA genotyping. Conservation Genetics 8: 475–488.

Deines AM, Bbole I, Katongo C, Feder JL, Lodge DM (2014) Hybridisation between native Oreochromis species and introduced Nile tilapia O. niloticus in the Kafue River, Zambia. African Journal of Aquatic Sciences 39: 23–34.

Eccles DH (1992) A Field Guide to the Freshwater Fishes of Tanzania, FAO Publications, Rome.

Lind CE, Brummett RE, Ponzoni RW (2012) Exploitation and conservation of fish genetic resources in Africa: issues and priorities for aquaculture development and research. Reviews in Aquaculture 4: 125–141.

Ndiwa TC, Nyingi DW, Agnèse JF (2014) An important natural genetic resource of *Oreochromis niloticus* (Linnaeus, 1758) threatened by aquaculture activities in Loboi drainage, Kenya. PloS One 9: e106972.

Petr T (1974) Hombolo Dam (Dodoma Region, Tanzania)- a brief account of the limnology with emphasis on the fish population. Makerere University. 29pp.

Seegers L (1996) The fishes of the Lake Rukwa drainage. Musee Royal de l’Afrique Central, Tervuren, Belgique, Sciences Zoologique 258: 1–396.

Shechonge A, Ngatunga BP, Bradbeer SJ, Day JJ, Freer JJ, Ford AGP, Kihedu J, Richmond T, Mzighani S, Smith AM, Sweke EA, Tamatamah A, Tyers AM, Turner GF, Genner MJ (2018a) Widespread colonization of Tanzanian catchments by introduced *Oreochromis* tilapia fishes: the legacy from decades of deliberate introduction. Hydrobiologia https://doi.org/10.1007/s10750-018-3597-9.

Shechonge A, Ngatunga BP, Tamatamah R, Bradbeer SJ, Harrington J, Ford AGP, Turner GF, Genner MJ (2018b) Losing cichlid fish biodiversity: genetic and morphological homogenization following colonisation by introduced species. Conservation Genetics 19: 1119–1209.

Shemsanga C, Muzuka ANN, Martz A, Komakech HC, Elisante E, Kisaka M, Ntuza C (2017) Origin and mechanisms of high salinity in Hombolo Dam and groundwater in Dodoma municipality Tanzania, revealed. Applied Water Science 7: 2883–2905.

Trewavas E (1983) Tilapiine fishes of the genera Sarotherodon, Oreochromis and Danakilia. British Museum (Natural History) Publications. 583 pp.

